# Sequence-dependent orientational coupling and electrostatic attraction in cation-mediated DNA-DNA interactions

**DOI:** 10.1101/2023.07.01.547339

**Authors:** Weiwei He, Xiangyun Qiu, Serdal Kirmizialtin

## Abstract

Condensation of DNA is vital for its biological functions and controlled nucleic acid assemblies. However, the mechanisms of DNA condensation are not fully understood due to the inability of experiments to access cation distributions and the complex interplay of energetic and entropic forces during assembly. By constructing free energy surfaces using exhaustive sampling, and detailed analysis of cation distributions, we elucidate the mechanism of DNA condensation in different salt conditions and with different DNA sequences. We found that DNA condensation is facilitated by the correlated dynamics of localized cations at the grooves of DNA helices. These dynamics are strongly dependent on salt conditions and DNA sequences. In the presence of magnesium ions, major groove binding facilitates attraction. In contrast, in the presence of poly-valent cations, minor groove binding serves to create charge patterns leading to condensation. Our findings present a novel advancement to the field and have broad implications for understanding and controlling nucleic acid complexes *in vivo* and *in vitro*.

## Introduction

Nucleic acids are the key molecular building blocks that store and convey genetic information. The precise higher-order structures of nucleic acids are often prerequisites for vital biological processes. A direct example of their organization in biological organisms is that meter-long DNA helices are compressed and confined inside the cell nucleus in the range of 10 ^−6^ meters.^1–4^ The process of compacting double-stranded (ds) DNA, the prevalent physical form of the genome, is technically known as DNA condensation, which is essential for gene regulation in all forms of life.^5^ DNA condensation is counterintuitive, as DNA strands are highly charged, suggesting an organized assembly of like-charged molecules.^6^ One critical component of macromolecular assemblies of nucleic acids is oppositely charged molecules, such as the histones in eukaryotic chromatin and the cationic lipids in lipid nanoparticles.^7–9^ Remarkably, a wide range of multivalent cations, such as naturally occurring polyamines spermidine^3+^, spermine^4+^ and inorganic cobalt hexamine (Co(NH_3_)_6_^3+^).^10–13^ Even divalent cations, have been observed to induce structured aggregates of DNA molecules *in vitro*.^14^ Despite extensive studies, the molecular mechanisms of DNA condensation, the subject of this study, remain under debate.^6,14,15^

Decades of research have yet to reach a consensus on the physical operating principles of the seemingly simple, nonetheless multifaceted, three-component system of DNA, ions, and solvent molecules. Competing theoretical models, such as the Kornyshev-Leikin (KL) zipper mechanism,^16^ the tightly bound ion ansatz,^17^ correlation effects,^10,18–20^ hydration formalism,^21^ and cation-bridging model, ^22^ have all succeeded in characterizing and explaining the spontaneous assembly of DNA molecules. However, each model usually addresses one pertinent aspect of the system and has a limited scope of applicability. For instance, the cation-bridging model can only be applied to biogenic polycations, such as spermine^4+^, but was not applicable to point-charge cations, such as divalent cations. In the KL mechanism, as another example, the binding of cations to dsDNA major grooves assumes a ‘static’ binding that is unrealistic due to the known cation density fluctuations. As a result, the model fails to predict DNA condensation by minor-groove binders or the recently reported DNA aggregation in alkaline-earth metal ions (Mg^2+^, Ca^2+^).^14^ In addition, while the hydration force proposal^23^ was based on experimental observation and theoretical formulation, its causal relationship with electrostatics is difficult to appraise.^24^

On the other hand, the challenges in arriving at a unified theory may be implicated in the diverse ranges of experimental conditions capable of condensing DNA. Consider the case of random-sequence dsDNA in divalent Mg^2+^ salts. Perturbing the solvent by adding ethanol condenses DNA; raising the cation valence with Co(NH_3_)_6_^3+^ or polyamines condenses DNA. To complicate it even more, chemical modifications of the ribose (e.g., 2’-OH addition to turn DNA to RNA) or the base (e.g., methylation of cytosine) have been shown to substantially weaken attraction.^22,25^ More recently, we showed that switching to homopolymeric sequences can also lead to DNA condensation.^14^

No existing analytical theories can describe all observations. Recent theoretical efforts have focused on using all-atom molecular dynamics (MD) simulations to dissect the different facets of DNA-DNA interactions.^6,22,26,27^ As a result, several distinctive molecular mechanisms have been identified, such as ion bridging with chain-like polycations,^20,22^ the constructive roles of externally-bound interfacial cations, and the destructive effects of deep helical grooves internalizing cations.^28,29^ Albeit powerful, MD simulations are subject to inaccuracies in the force fields and difficulties in sampling the multiple time and length scales of the macromolecular system. These challenges have prevented MD simulations from providing a holistic view of all relevant degrees of freedom, especially the configurational space of DNA helices.

To address such challenges, we have investigated the utilities of advanced exhaustive sampling techniques and refined empirical potentials in all-atom molecular simulations of DNA-DNA interactions. Our computational approach enabled us to develop a reliable theoretical protocol for studying nucleic acid interactions across various conditions. We were able to accurately reproduce DNA condensation/dissolution experiments under different conditions and sequences. Expanding upon previous studies that exclusively utilized inter-helical spacing as the reaction coordinate for studying DNA-DNA interactions, we included azimuthal angle as an additional variable. This added complexity has yielded novel findings that shed light on the mechanism of DNA condensation. Our simulations indicate that the azimuthal angle plays a crucial role in positioning the helices to facilitate attraction. To investigate the mechanism of like-charge attraction, we compared the recently observed divalent-mediated dsDNA attraction with the well-known polyvalent-induced DNA condensation. We examined the effect of DNA sequence on DNA-DNA interaction by contrasting a repeating poly(A)-poly(T) sequence (ATDNA) with a random sequence (MixDNA) that mimics genomic DNA. In addition, to investigate the role of cations in modulating DNA-DNA interactions, we studied systems with pure divalent magnesium salt, as well as ionic mixtures of sodium/magnesium and sodium/magnesium/spermine.

Through our atomistic simulations, we were able to decompose the free energy landscape into energetic and entropic factors, providing valuable insights into the underlying forces driving DNA condensation. We have observed that cation distribution is the primary factor contributing to DNA-DNA interactions, with both energetic and entropic components playing a role in modulating these interactions. In homopolymeric ATDNA sequences, enhanced cation binding and dynamics in the major groove of DNA significantly contribute to DNA-DNA attraction induced by Mg^2+^. Although DNA condensation is an overall process of entropy loss for cations, we have observed that the loss of cation entropy is dependent on the DNA sequence. Specifically, ATDNA pairs exhibit a lower entropy loss than mixed DNA sequences due to the enhanced dynamics of groove-bound cations. These cations play a critical role in DNA-DNA attraction and inter-helical orientational coupling through charge-charge correlations. In a Na/Mg mixture solvent, we observed that attraction forces weakened due to ion competition, leading to inadequate electrostatic screening. However, the addition of polyvalent cations, such as spermine^4+^, resulted in a shift in the equilibrium towards attraction due to the specific binding of the polyvalents to the minor grooves. In contrast to divalent-induced DNA condensation, the attraction induced by spermine was caused by chain-like cations bridging adjacent phosphate groups by utilizing minor grooves.

Our results demonstrate excellent qualitative and quantitative agreement with experiments and reveal a distinctive mechanism of DNA condensation. The regular, extended charge patterns in the grooves create a salt-dependent “dynamic chain” of mobile cations, resulting in inter-helical orientational coupling and azimuthal ordering. We propose that surface structure and sequence-dependent binding of cations represent a novel principle for DNA interactions. Our findings have implications in genome packing, molecular recognition, and molecular assembly.

## Theory and Methods

All-atom molecular dynamics simulations were used to study the interactions between pairs of double-stranded DNA (dsDNA). Two DNA sequences were investigated: a 20-bp duplex consisting of two homopolymeric chains (dA_20_ and its complementing strand), referred to as ATDNA; and a quasi-random sequence (GCA TCT GGGC TATA AAA GGG and its complement), referred to as MixDNA. We positioned each DNA pair in parallel (see Fig. 1a) and sampled conformations of distances and orientations. To investigate the solvent dependence of DNA-DNA interactions, we studied pure Mg, Mg/Na, and Mg/Na/Spermine mixtures. Well-tempered metadynamics simulations allowed us to construct free energy landscapes.^30^ We analyzed the electrostatic and entropic contributions of DNA and its partners, as well as hydration. Details of the simulation approach, solution conditions, and analysis methods are summarized below and explained in depth in the Supplemental Materials and Methods.

**Figure 1:**
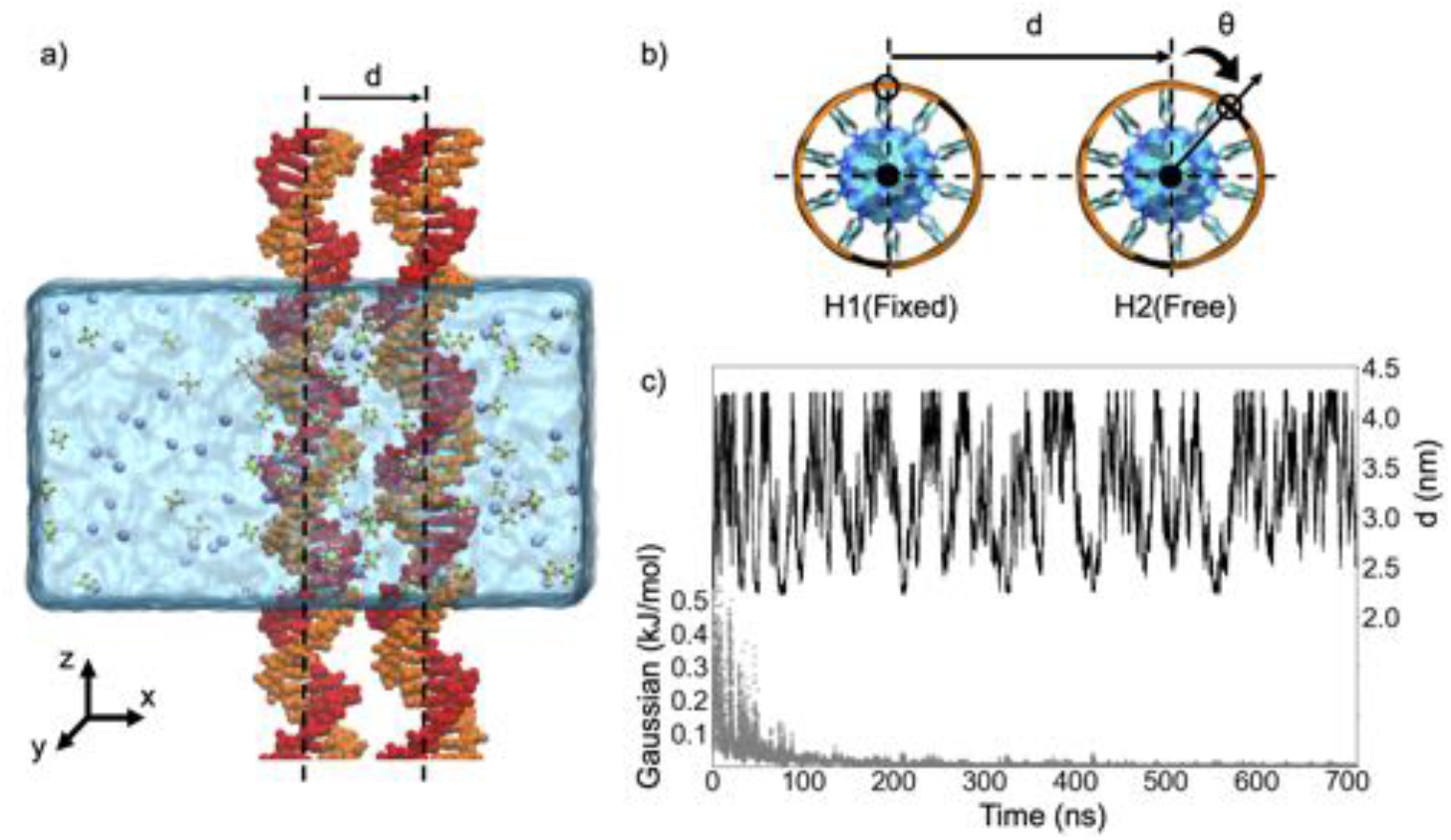
Simulation setup of the parallel DNA duplexes. a) Simulation box with explicit water and ions. Periodic boundary conditions along the z-axis allowed extending the DNA to infinite length, mimicking DNA-arrays. b) The reaction coordinate of DNA-DNA interactions is described by inter-helical distance *d* measured in (*x, y*) plane, and *θ*, the rotation of helix 2 (H2) with respect to the helix (H1) restrained in orientation. To sample conformations in *d, θ*, Well-tempered metadynamics (WTMD) simulations is employed. c) The convergence of sampling monitored by the time evolution of the Gaussian height (gray) and inter-helix spacing *d* (black).

### Molecular modeling and simulation setup

Both dsDNAs were built in B-form using Nucleic Acid Builder (NAB).^31^ DNA pairs were placed in a simulation box of 11.8 ×11.8 × 6.8 *nm*^3^. We aligned them along their long axis such that the DNAs extend to infinity under periodic boundary conditions, mimicking long DNA arrays used in experiments. Simulations were carried out using the GROMACS 2018.5^32^ suite of programs. We used the amber99sb parmbsc0^33–36^ force field to represent DNA, water was modeled by TIP3P.^37^ For Mg, Na, and Cl we used NBFIX^26^ parameters. For spermine we used general Amber parameters with partial charges derived from Hartree Fock (HF) calculations that were fitted using the RESP approach.^38^ Further details of the modeling and simulation methods are explained in the Supplementary Materials.

### Well-tempered metadynamics

To exhaustively sample the conformational space of DNA pairs, we performed Well-tempered metadynamics implemented in PLUMED.^30,39^ We reduced the dynamics to two straightforward collective variables (CV): the inter-helical distance (*d*) and the axial rotations of helices relative to one another (Fig. 1a-b). For ease of visualization and analysis, we constrained the rotations and translations of one helix (H1) using the enforced rotation implementation in GROMACS,^40^ while allowing the other helix (H2) to freely move. The fluctuations of the collective variables were monitored and the convergence of the simulations was assessed by evaluating the amplitudes of the Gaussian heights deposited in the space of the CVs (Fig. 1c). WTMD sampling was considered converged when the Gaussian hills decayed and reached asymptotic values. The block analysis was used to further examine the convergence (Fig. S3). Detailed descriptions of the WTMD setup are given in the Supplementary Materials. Details of the WTMD theory and our application to DNA-DNA interactions can be found elsewhere.^30,41^

### Data analysis

From the particle positions in a simulation box of volume *V*, we compute the average cation number density

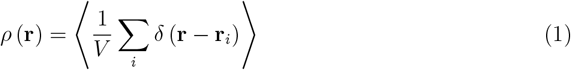

where r ≡ (x,y,z) represents an arbitrary point in space. ⟨. ⟩ represents the ensemble average obtained from a 300-ns brute-force MD simulation. We use a spatial resolution of 1Å.

Due to the helical nature of the duplex, we transform the density profile to the cylindrical coordinates:

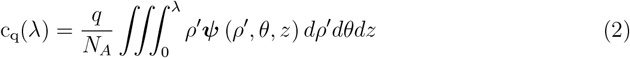

where λ represents the distance from the center of the helix, *N*_*A*_ is the Avogadro constant and *q* is the valence of the specified ion. Based on the cylindrical geometry, we compute the average number of excess cations condensed on the DNA up to a distance *R*:

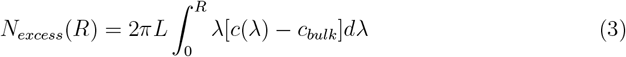

where *c*(λ) is the average cation concentration around a cylindrical coordinate at radial distance λ, *c*_*bulk*_ is the bulk concentration of the cation from the center, and *L* is the length of the DNA long axis.

Similarly, for each point in space, we computed the charge density *ρ*_*q*_(**r**). We probe the electrostatic potential acting at an arbitrary point *r*_1_

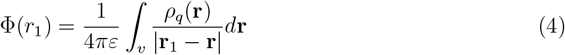

where the integral is evaluated over a sub-volume, *v* forming a shell of 10 Å thickness on the DNA’s surface. The cut-off is determined based on the Debye length at the salt condition. From Φ(*r*), we computed the stored electrostatic potential energy between helices at an arbitrary inter-helical distance, *d*, as:

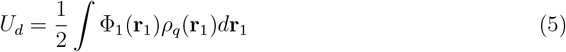

To accurately assess the contribution of thermodynamic potentials, we also computed the entropy change upon assembly. For that, we divided the entropic contributions into the conformational entropy of DNA pairs, and the entropy of the solvent (cations and water). The first one is estimated using the multiscale-cell-correlation (MCC) theory,^42,43^ where the total entropy is the sum of the following:

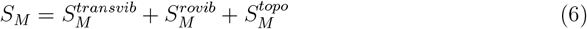

Here 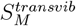 and 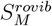 are translational and rotational vibrations of macromolecule, while 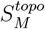 accounts for the topographical entropy. Due to the well-defined structure of the double helix, we found the changes of 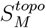 term negligible.

To estimate the entropy of Mg^2+^ ions and water, we employed the two-phase thermodynamic (2PT) method.^44–47^ In this method, the density-of-state (DoS) is represented by solid-like (*S*^*s*^(*v*)) and gas-like (*S*^*g*^(*v*)) components, *S*(*v*) = *S*^*g*^(*v*)+ *S*^*s*^(*v*), and the entropy is estimated from the DoS associated with the atomic velocities (*v*) decomposed into contributions from molecular translation (*S*_*trans*_(*v*)), rotation (*S*_*rot*_(*v*)), and vibration (*S*_*vib*_(*v*))^46^:

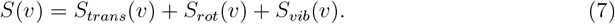

Details of the theory can be found in Ref,^44^ and our implementation of the method is detailed in the Supplementary.

To determine the thickness of the hydration layer, we computed the tetrahedral order parameter:^48^

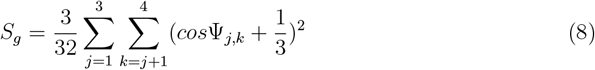

where Ψ_*j,k*_ is the angle subtended at the central atom between the *j*-th and *k*-th bonds. The factor of 3/32 serve to adjust *S*_*g*_ in the range 0 < *S*_*g*_ < 1. We binned the data into cylindrical shells with a bin size of 0.5 Å.

## Results

Due to the high persistence length (150 bp), DNA-DNA interactions can be modeled as parallel rods. As these interactions are additive, the forces between the long DNA strands increase in strength, resulting in a strong driving force at the macroscopic level, although relatively weak at the base-pair level. Achieving accurate computational modeling of these interactions requires capturing a free energy change of 0.01 to 0.1 kJ/mol/bp, which is challenging due to inaccuracies in force fields and the rugged energy landscapes that must be sampled. However, by extensively sampling inter-DNA distances and orientations using a force field refined based on solution studies of DNA interactions, we were able to successfully replicate experimentally consistent DNA interactions, enabling us to conduct a comprehensive analysis of cation-mediated interactions.

### WTMD establishes sequence-dependent DNA-DNA attraction in Mg^2+^

Using our approach, we first looked at the ATDNA, the homopolymeric DNA sequence that shows condensation in experiments.^14^ We compared the ATDNA sequence with a sequence mimicking the genomic DNA, constructed by introducing an about equal composition of each nucleobase, abbreviated as MixDNA henceforth. Both sequences are studied under the same salt condition to provide a fair comparison. The free energy profiles of the two sequences are shown in Fig. 2a-d. The free energy projected on inter-helical distances suggests that the ATDNA pair has the energy minimum at *d ∼* 2.8 nm (Fig. 2a), consistent with the equilibrium inter-helical distance measured by experiments.^14^ The free energy of binding Δ*F* = *F*(2.8 *nm*) − *F*(4.0 *nm*) ≈ (−0.15) kJ/mol/bp agrees well with the estimated value from osmotic stress measurements.^49^ This difference in energy between the bound and unbound states results in ≈ (45 − 75) kJ/mol of stabilization for DNA pairs of 300-500 bp, which is sufficient to overcome the thermal energy, leading to condensation, as observed.^14^

**Figure 2:**
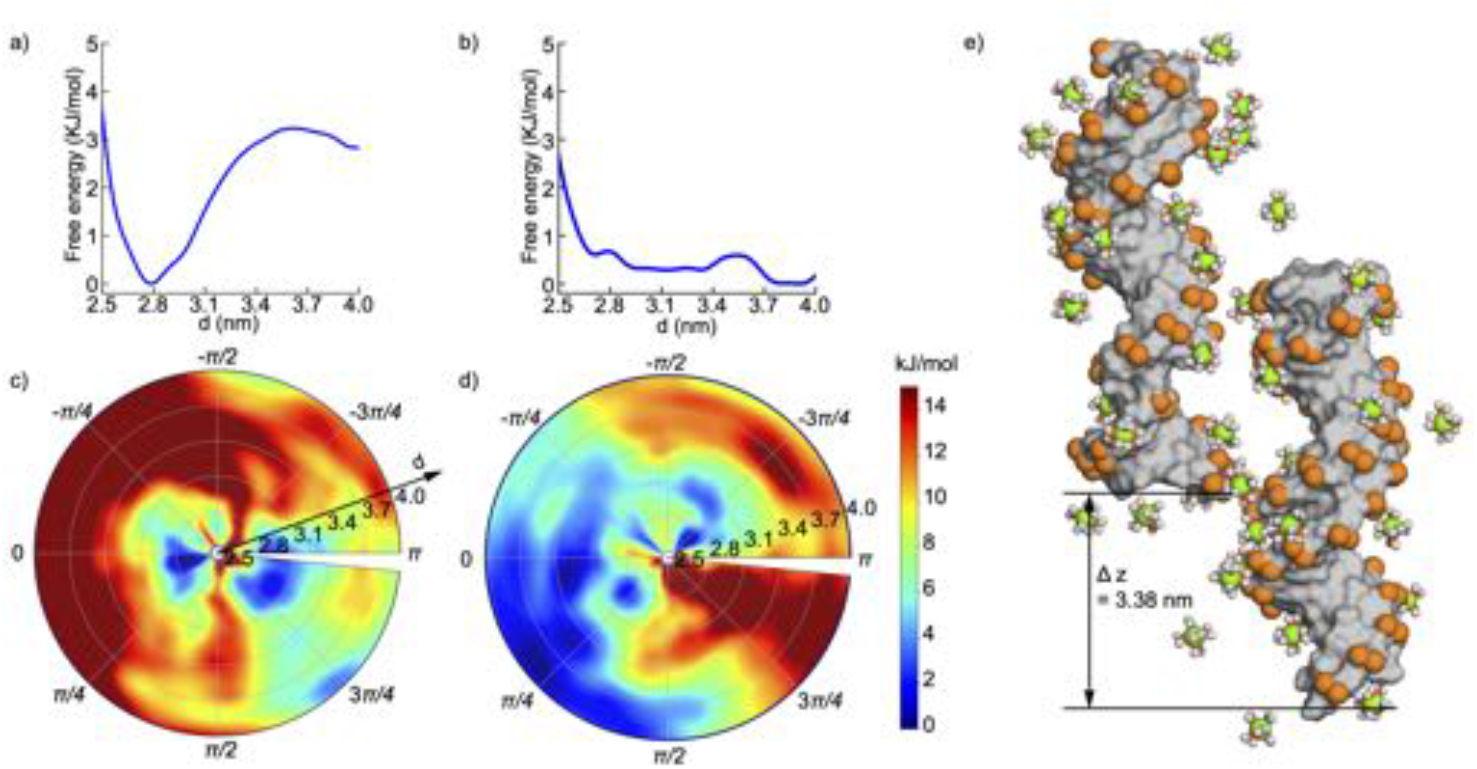
Free energy profile of sequences ATDNA and MixDNA in pure Mg^**2+**^. a) The free energy profile of ATDNA as a function of inter-helical spacing *d*. b) The free energy profile of MixDNA. c-d) The free energy surface in two dimensional (*d, θ*) polar chart. In the polar plot of the free energy surface, the radial distance from the origin represents the inter-spacing distance (arrow), denoted as *d* in panels a and b, while the angular coordinate corresponds to the azimuthal rotation *θ*. c) ATDNA and d) MixDNA. e) The representative conformation of ATDNA at the energy minimum, (*d ∼*2.77 nm, *θ* 0.25 rad), See Movie 1 in SI for the dynamics.

While the free energy profile of ATDNA shows attraction, the free energy of the MixDNA lacks any deep minimum at short DNA-DNA distances (Fig. 2b). The downhill nature of the energy profile of MixDNA suggests spontaneous dissociation of DNA arrays. This observation is in accord with experiments of the genomic DNA.^14^

To provide further details on the DNA-DNA interactions, we present the free energy landscape in two dimensions (Fig. 2c-d). The polar plots display energy values in interhelical distance, *d*, and azimuthal angle, *θ*. Consistent with the pictures in Fig. 2a-b, the FESs show sequence-dependent behavior with angle correlations in ATDNA for shorter interhelical distances. The orientational coupling observed minimizes the distances between the phosphate backbone and the major grooves (Fig. 2e). In sharp contrast, MixDNA shows weak orientational coupling at short distances (Fig. 2c,d), a wide but visible correlation for MixDNA is also evident in longer distances (Fig. S1).

### Solvation shells show subtle differences between ATDNA and MixDNA

ATDNA duplexes exhibit attraction under the same conditions where MixDNA exhibits repulsion. To investigate the sequence-dependent causal factors, we analyze and compare relevant thermodynamic quantities of DNA, cations, and water in detail. Given the critical role of solvent, we examine the changes in water and cation coordination in addition to the electrostatic energy stored in the solvation shell (Fig. 3).

**Figure 3:**
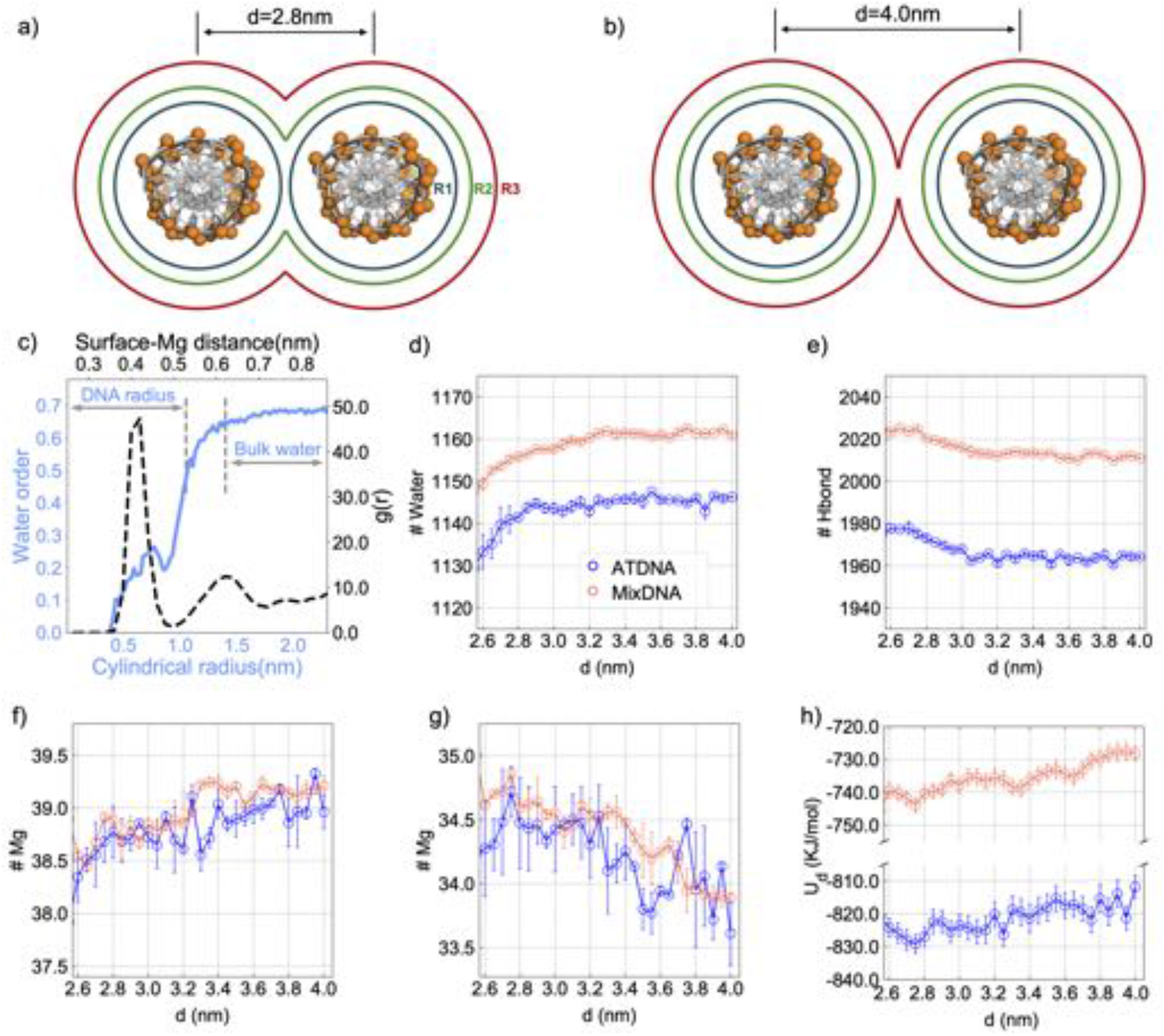
Solvation shell dynamics in free to condensed phase transition. Graphical representation of the different layers of DNA solvation shell comprised of 3.5A hydration (R1), 6.0A tight cation binding (R2) and 10.0A of Debye layer (R3) respectively. a) condensed phase solvation layers and b) free state. c) Tetrahedral water (blue) and radial distribution function (dashed black) analysis of Mg^2+^ around the surface of a DNA helix. d) Change in bound water molecules in R1 as a function of inter-spacing *d*. e) Change in number of hydrogen bonds within R1 as a function of inter-spacing *d*. f) Change in Mg^2+^ in the outer layer (R3), g) Number of bound Mg^2+^ ions, (R2 region). h) Change of stored electrostatic energy (*U*_*d*_) within the R3 layer as a function of inter-DNA spacing.

To facilitate analysis, we have divided the surface of the DNA into three regions (Fig. 3a). The first region, denoted as R1, corresponds to the bound water region identified by tetrahedral water analysis (Fig. 3c, left y axis), and spans from 0 to 3.5 Å. The second region corresponds to the bound cation region identified by RDF and spans from 0 to 6 Å(Fig. 3c, right y axis). The third region is defined based on the Debye length at our salt condition and spans from 0 to 10 Å. We define the DNA pairs as a condensed state when the inter-helical distance is at 2.8 nm, and the free state is defined when pairs are at 4.0 nm (Fig. 3a-b). We monitored the changes in the solvation shell as the double strands transitioned from the free state to the condensed phase along the condensation path. The path was constructed using 500 snapshots from inter-helical distances, equally spaced with a resolution of *δd* = 0.5 Å.

We observed that MixDNA has a stronger hydration shell compared to ATDNA (Fig. 3d-e).The coordination number of bound water (at R1) shows a reduction as the DNAs come closer, suggesting water release from the DNA surface. Interestingly, despite the reduction of water from the solvation shell of R1, the number of hydrogen bonds shows an increase as the two DNA pairs approach one another, suggesting water bridging between the two DNA hydration layers. MixDNA, which has a denser water shell, has a greater number of water molecules and hence a higher number of hydrogen bonds throughout the pathway. The differences between ATDNA and MixDNA remain consistent as the DNAs transition from the free to the condensed phase.

The coordination number of cations in the R3 region shows a reduction upon DNA condensation, while the coordination of cations to the R2 region increases (as shown in Fig. 3f-g), indicating a reorganization of cations in the outer and inner solvation shell as the DNA pairs come closer. However, despite these changes, the number of counterions does not exhibit a strong sequence dependence to explain the distinct behavior observed in the free energy landscape.

Although there are substantial similarities in the cation coordination between the two sequences, we have observed a notable difference in the stored electrostatic energy, *U*_*d*_, (as described in the methods) between ATDNA and MixDNA (as shown in Fig. 3h)). Notably, the magnesium distribution around ATDNA creates a better screening for the double helices compared to MixDNA leading to lower *U*_*d*_ values throughout the path.

To understand the structural differences between the cation distributions that lead to differences in the electrostatic response of the duplexes, we have plotted the ion densities in the form of radial distribution and 3D number densities (as shown in Fig. S2). For both sequences, we have observed a two-layer ion distribution around DNA. However, the localization patterns of cations around the two layers are different in the ATDNA and MixDNA duplexes. In the case of ATDNA, a uniform and well-defined distribution of cations is observed in the major groove. On the other hand, in the case of MixDNA, discrete binding pockets reflecting the heterogeneity of the sequence are evident. The cation distributions of the DNA surface show stronger DNA-Mg interactions in ATDNA (as shown in Fig. S2), indicating that the ATDNA sequence provides a stronger electrostatic field to attract cations. The difference between ATDNA and MixDNA is that in the ATDNA, the major groove surface is decorated with two highly electronegative sites forming charge patterns that do not exist in the case of MixDNA^14^. Additionally, ATDNA possesses a wider major groove^50^, which results in a shorter phosphate-to-phosphate distance between the two strands, leading to a higher negative charge density at the surface of the DNA. The higher peak intensity in the radial distribution function (RDF) (as shown in Fig. S2) observed in the case of ATDNA highlights the importance of the inter-chain phosphate group distances.

The electrostatic interaction in the solvation shell favors DNA condensation in both sequences, but ATDNA provides better enthalpic stabilization on the absolute scale, yet relative energy changes remain similar. However, the question remains regarding how the entropic factors compare between the two sequences. To assess the entropic factors, we have partitioned the total entropy in the solvation shell (as defined above) into the components of cation, water, and DNA. We define the change in entropy as *ΔS* = *ΔS*_*cation*_ + *ΔS*_*DNA*_ + *ΔS*_*water*_, where *ΔS*_*x*_ = *S*_*x*_(2.8*nm*) − *S*_*x*_(4.0*nm*) represents the entropy change for each term, with *x* =*total, cation, DNA*. The details of computing each entropy term are explained in the Methods section and the SI. Our findings are reported in Table 1.

**Table 1:** Entropy change during the transition

| | $-T\Delta S_{Mg^{2+}}$<br>(kJ/mol) | $-T\Delta S_{DNA}$<br>(kJ/mol) | $-T\Delta S_{water}$<br>(kJ/mol) | $-T\Delta S_{total}$<br>(kJ/mol) |
| --- | --- | --- | --- | --- |
| ATDNA | 1.87±2.10 | 0.62±0.23 | -13.04±1.13 | -10.55±2.40 |
| MixDNA | 7.04±1.36 | 0.83±0.54 | -14.11±1.57 | -6.24±2.15 |

Based on our analysis, we found that the displacement of water makes a significant contribution to the entropy and overall stability of the condensed phase (as shown in Table 1). In contrast, the entropy of DNA plays a minor role. Interestingly, we have discovered that cation entropy shows a notable difference between condensed and free states. The most significant difference between the two sequences is their cation entropy changes; an entropic penalty 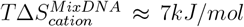 makes DNA condensation less likely for MixDNA. ATDNA’s unique topology, with two continuous binding layers and a wider major groove that facilitates dynamic exchanges of condensed cations^41^ lead to an overall free energy penalty of 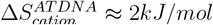. As a result, ATDNA offers a more stable condensate compared to MixDNA at the same salt condition.

### Distinct azimuthal ordering of ATDNA in ion-mixtures

The surface structure of ATDNA results in lower electrostatic energy and higher mobility of cations. This property stabilizes DNA pairs in the presence of pure MgCl_2_ solution. The unique ability of ATDNA to orient itself and self-assemble in divalent cations raises the question of whether other cations, especially those present in physiological conditions, can mediate attraction and azimuthal ordering in a similar way. Physiological conditions for DNA consist of cationic mixtures ranging from simple ions such as Na^+^/K^+^ and Mg^2+^ to polyelectrolytes, including polyamines and oligopeptides. To answer that we investigate two mixture conditions: 1) a mixture of Mg^2+^/Na^+^, denoted as Na-Mg henceforth; and 2) a mixed solution of Mg^2+^/Na^+^/Spermine^4+^, denoted as Na-Mg-Sp. The concentrations of each species in the mixture were adjusted to maintain a constant total Cl^−^ concentration in the simulation box. Details of the molecular setup of the mixtures can be found in the Supplementary Information. Similar to the pure Mg^2+^ solution, we used WTMD to construct the FES of the duplex pairs.

The free energy landscape of the two ionic mixtures is presented in Fig.4a-b. In Na-Mg, we observe a repulsive downhill potential where the lower energy states correspond to the duplex pairs being far apart. This observation is reminiscent of MixDNA in pure MgCl_2_. Similar to MixDNA in MgCl_2_, no azimuthal correlations are apparent from the FES as shown in Fig.4c.

**Figure 4:**
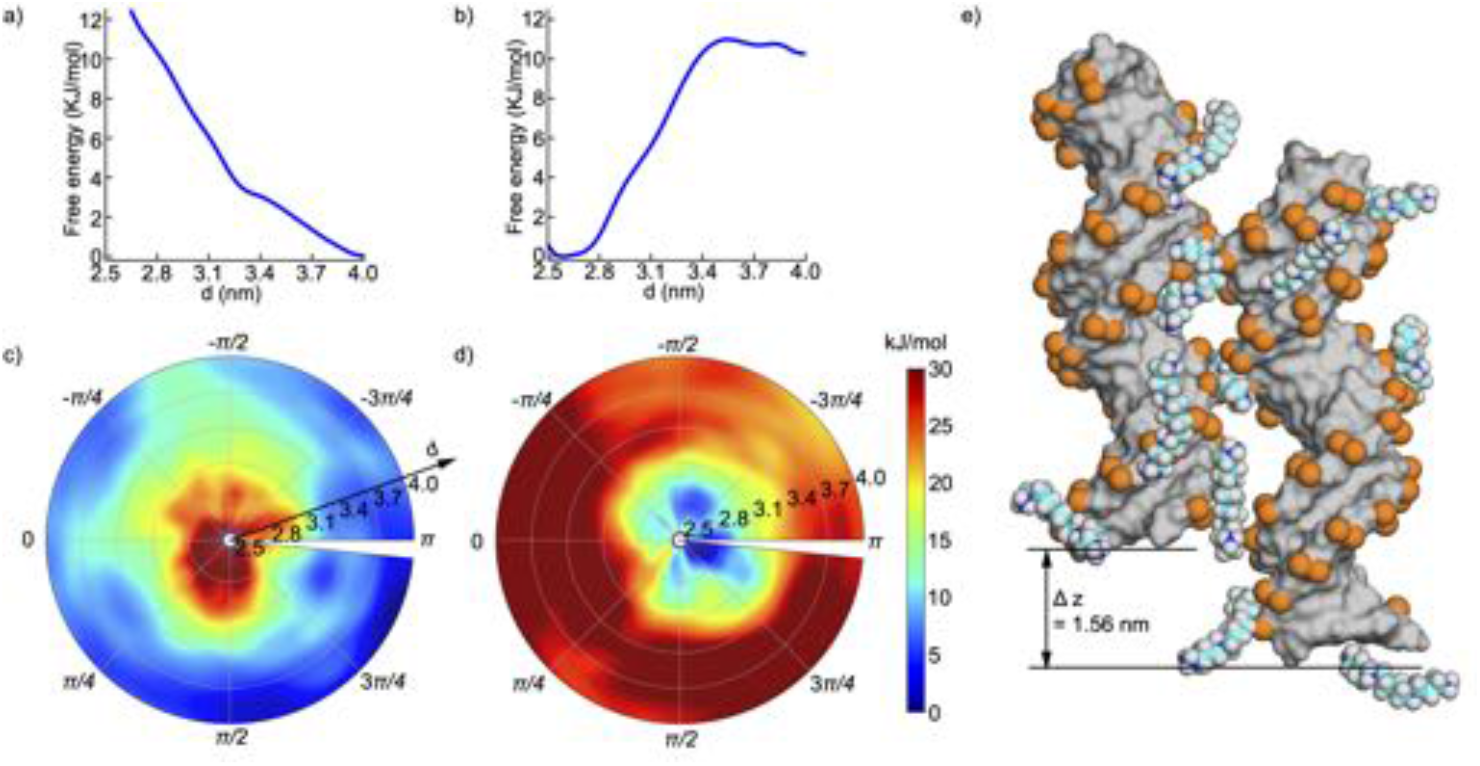
Free energy surface sampled by metadynamics simulation projected onto the two collective variables (*d, θ*) for ATDNA in physiological salt conditions. a) The free energy profile along the reaction coordinate *d* for ATDNA in Mg^2+^/Na^+^, showing monotonic descending order. b) The free energy profile along the reaction coordinate *d* for ATDNA in Mg^2+^/Na^+^/Spermine^4+^. c-d) The 2D FES map inside polar chart projected on (*d θ*) c) for ATDNA in Mg^2+^/Na^+^ and for d) ATDNA in Mg^2+^/Na^+^/Spermine^4+^, respectively. e) The snapshot shows the representative conformation of ATDNA in Mg^2+^/Na^+^/Spermine^4+^ at the energy minimum, (*d ∼* 2.59 nm, *θ ∼* 2.51 rad), and illustrates the Spermine^4+^ ligand bridging between DNA pairs. (See the Movie 2 in SI)

Unlike the Na-Mg solution, the FES of Na-Mg-Sp exhibits a distinct minimum at close inter-helical distances (Fig.4a-b). This observation is not surprising since Spermine^4+^ is an effective condensing agent that can condense both long and short DNA duplexes, regardless of whether they are natural or synthetic.^51^ Our results demonstrate a global minimum at *d ∼* 2.6nm (Fig.4d), which is in good agreement with previous simulation and experimental studies as highlighted in Ref,^22^ In Na-Mg-Sp, similar to pure MgCl_2_, we observe a well-defined angular preference with a deeper free energy minimum than in pure MgCl_2_, albeit at very different azimuthal angles (2.51 vs. 0.25 rad for Na-Mg-Sp vs. MgCl_2_).

### Ion competition modulates DNA interactions in mixtures

To understand the molecular details underlying the different behaviors observed in the various salt conditions, we analyzed the average charge densities around the cylindrical coordinate of the H1 duplex in the free state, i.e., when the two duplexes are far apart. We computed the excess cation charge around the DNA helix from the concentration profiles. The density profiles and cumulative charges under the three salt conditions are shown in Fig. 5a-b.

**Figure 5:**
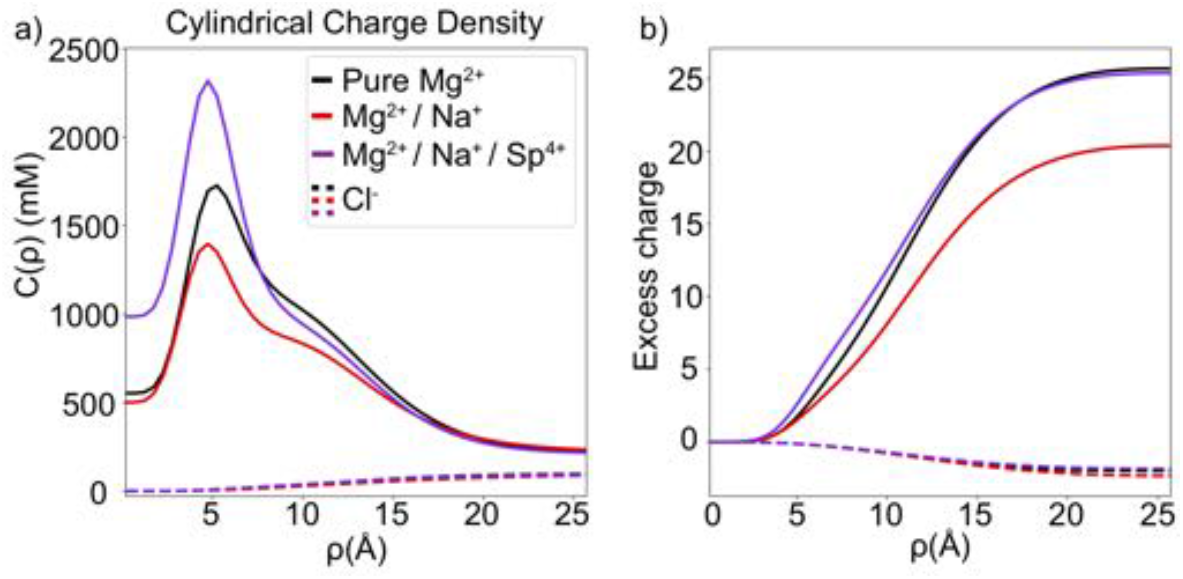
The comparison of cation charge distribution around the cylindrical axis of the DNA in different salt conditions. a) Concentration profile of the total charge as a function of distance from the center of DNA, *ρ*. b) The excess positive cation charge (solid) and negative charge (dashed)

The analysis of cation charge density reveals a critical difference between Na-Mg, which shows repulsion, and pure Mg^2+^ and Na-Mg-Sp, which show attraction. The common feature of the attraction cases is that the cation charge density is higher, resulting in an excess charge accumulation that effectively screens the negatively charged DNA charges (Fig. 5a, purple solid line). In the case of repulsion, the cation charge density is lower, resulting in poor screening of the DNA charges. These marked differences in overall DNA charge neutralization are consistent with the observed attractive and repulsive interactions.

To understand the reasons behind the differences in screening, we constructed three-dimensional ion density maps. We visualized the spatial distribution of cations around the DNA under each mixture condition (Fig.6a-c). As a reference point, we compared the cation distributions to pure Mg^2+^ (Fig. 6a). To obtain a more quantitative picture, we also computed the cations’ surface distribution function (SDF) (Fig. 7) by probing the cations from the surface of the DNA rather than using a spherical volume element as used in RDF (Fig. 7).

**Figure 6:**
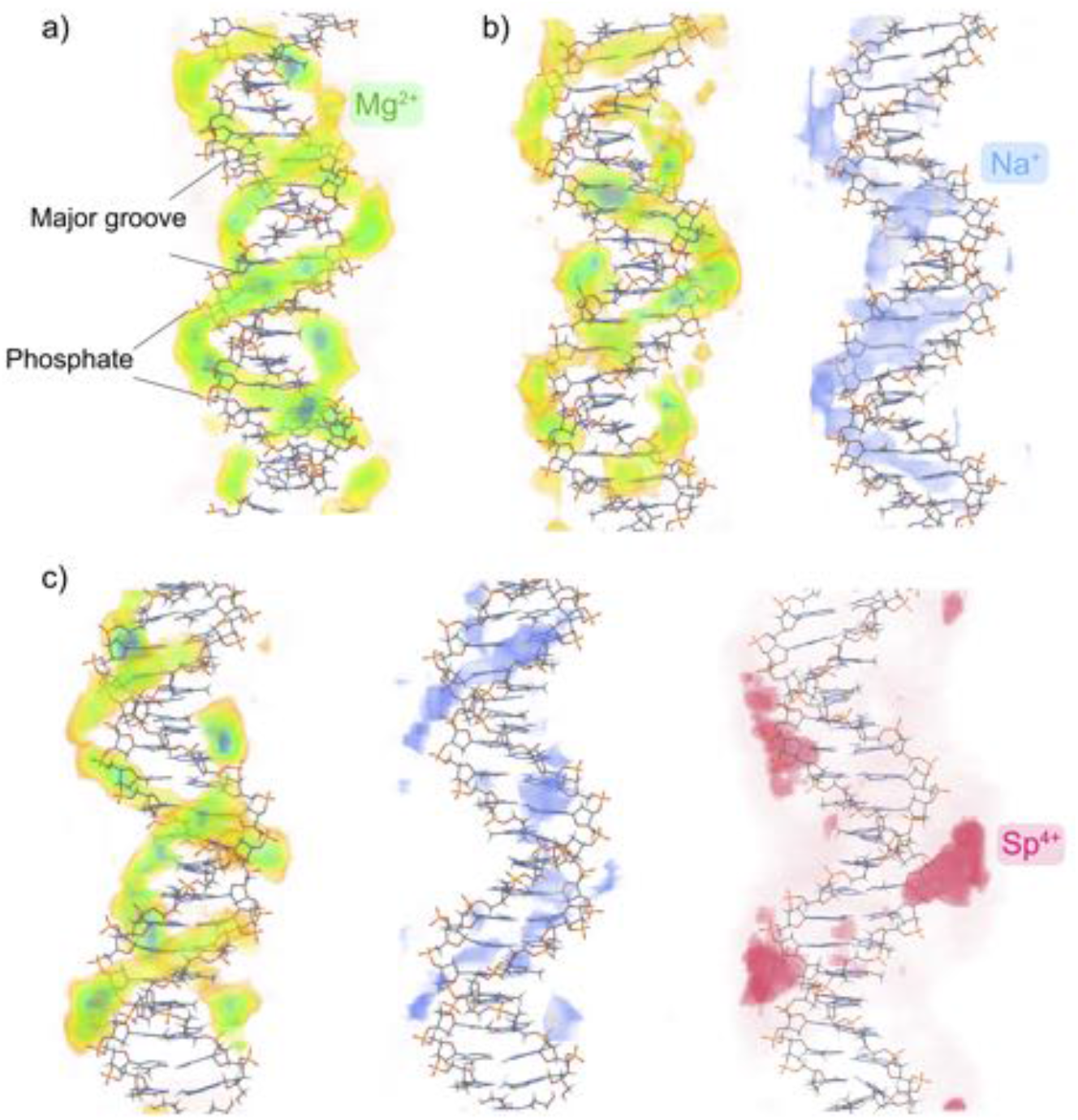
Spatial distribution of cations around the DNA duplex in different salt conditions. a) The 3D density map of cations in pure Mg^2+^ b) in mixtures of Mg^2+^ (green) and Na^+^ (blue), c) in mixtures of Mg^2+^ (green), Na^+^ (blue) and Sp^4+^ (pink).

**Figure 7:**
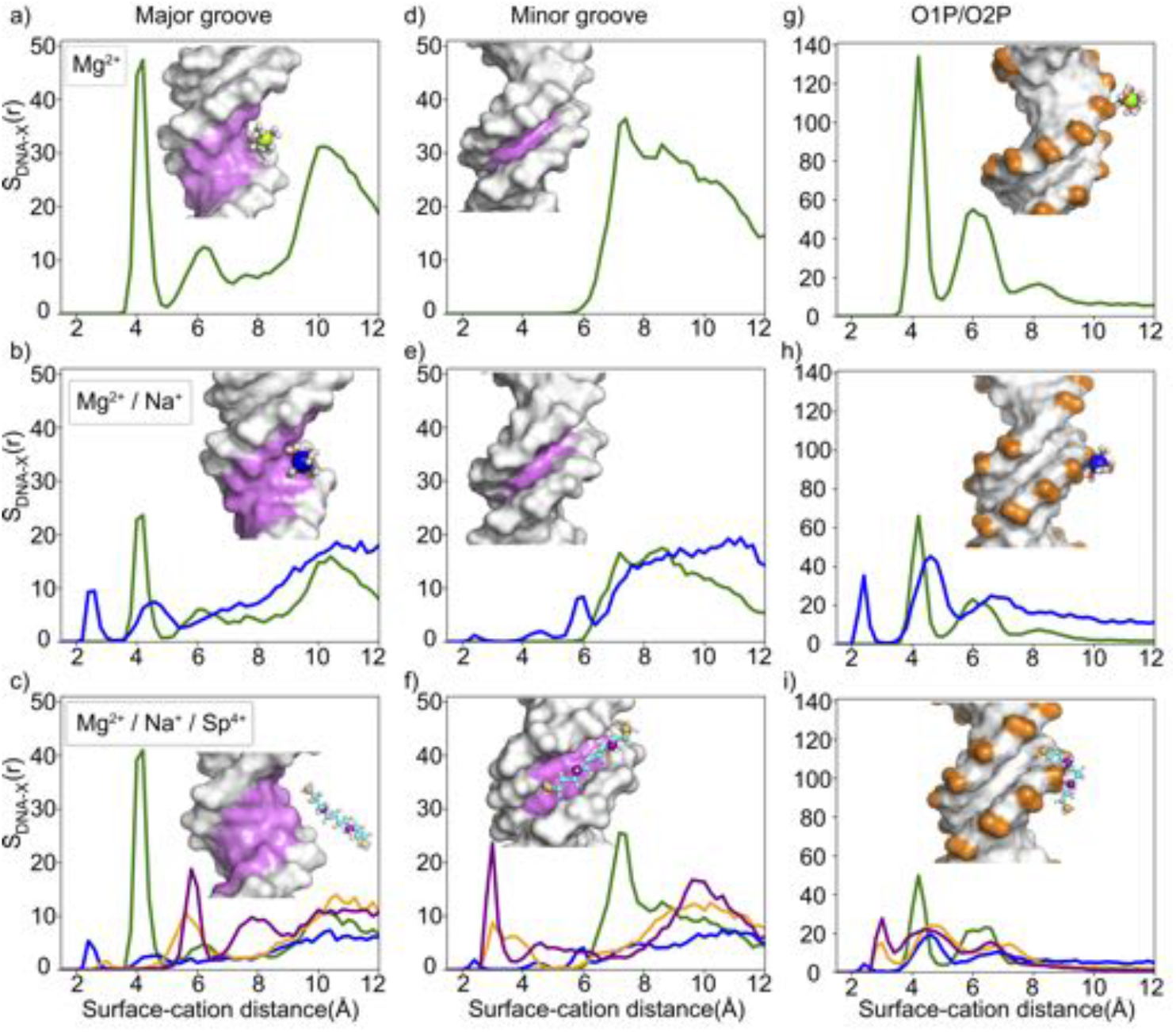
The comparison of the surface radial distribution function of cations around the DNA duplex in different salt conditions. a-c) major group atoms, d-f) minor groove atoms, and g-i) phosphate backbone. The x-axis represents the distance between cations and the surface of the selected group. For Spermine^4+^, we divided it into two groups as shown in the inset of c): terminal nitrogen atoms (orange) and non-terminal ones (purple).

A quick glance at the density maps reveals that cations compete with one another for the two DNA sites (major and phosphate backbone). The addition of Na^+^ to the Mg^2+^ salt impacts the two-layer binding of magnesium ions. We observe a weakening of Mg binding to the DNA surface. Indeed, the peak intensity of the SDF plots of Mg^2+^ ion in pure Mg and in Na-Mg mixture also shows this trend. Both the first layer and second layer of Mg binding to the DNA surface are impacted by the competition. The weakening occurs in the major groove and the phosphate backbone synonymously. The minor groove, which does not show any Mg binding, remains the same.

How a monovalent ion competes with a divalent ion remains an interesting question. A careful look at the SDFs shows that Mg^2+^ ions keep a fair distance away from the DNA surface, e.g., peaking at *∼*4.2*Å* distance compared with *∼*2.4*Å* surface distance for Na^+^ (see Fig. 7a-i). This can be attributed to the strong hydration shell of Mg^2+^, which remains largely intact when bound to DNA. Unlike Mg^2+^, the Na^+^ ions have a weaker solvation shell, leading to frequent dehydration events and almost equal Coulombic forces between hexahydrated divalent magnesium cations and dehydrated monovalent sodium cations.

Surprisingly, adding Spermine^4+^ to the mixture helps to partially regain most of the lost territories of the magnesium binding sites at the major groove. This observation is better seen when we look at the SDF (Fig. 7c in comparison to 7b). Indeed, Mg distribution shows a stronger peak in the Mg-Na-Sp mixture in comparison to Mg-Na; despite the latter having a higher concentration of magnesium cations. To achieve that, the Spermine^4+^ ions expel Na^+^ ions from the major and phosphate backbone regions (Fig. 7 c, i vs b and h). Unlike Mg, spermine acts as a competitor to sodium cations at all binding locations. Another notable observation is that, due to its linear shape and weak hydration shell, Spermine^4+^ affords to explore an alternative binding site. It mainly localizes around minor grooves (Fig. 4e, Fig. 6c, and Fig. 7f) leading to the two well-defined “chains” of cation densities along the helical geometry of ATDNA (Fig.6a).

Consistent with previous studies of DNA interactions in spermine,^22,27^ our simulations identify spermine’s preferential localization at the minor groove regions. As illustrated in Fig. 4e, the bridging effect is likely facilitated by its chain-like shape, capable of physically spanning the inter-helical space, leading to complex or disordered bridging configurations. Differently, our WTMD simulations that traverse inter-helical distances and azimuthal angles suggest measurable azimuthal coupling for ATDNA construct in the presence of spermine. The orientational coupling common for Mg^2+^ and spermine offers a more general mechanism for DNA-DNA interaction based on the correlated charges localized on the DNA surface. Based on our observation, we propose a zipper-like mechanism for like-charge attraction as a common mechanism for DNA-DNA interactions.

## Discussion and Conclusions

Motivated by the elusive physical principles underlying ion-modulated DNA-DNA interactions, we conducted a computational study of the recently observed sequence-dependent attraction between two different DNA constructs under various solution conditions. Using Well-tempered metadynamics and state-of-the-art analytical techniques, we provide unprecedented details on the configurational degrees of freedom and the thermodynamic contributions of all constituents. Importantly, comparing ATDNA and MixDNA offers a unique opportunity to probe the roles of DNA surface features, leading to the revelation of the unusual capacity of mobile, major-groove-bound divalent cations in mediating sequence-dependent DNA-DNA attraction.

Our study provides robust evidence for continuous cation distributions in the ATDNA major groove and the positioning of groove-bound cations next to the opposing phosphate backbone of the other helix, enabled by inter-helical orientational coupling. In contrast, MixDNA displays patched cation 3D distributions that do not register with opposing phosphate backbones in either the free or assembled states. As a result, the favorable electrostatic energy of ATDNA arises from a combination of stronger major-groove binding of Mg^2+^, delocalization of bound cations, and inter-helical azimuthal ordering, which collaborate to enhance one another. At a more fundamental level, these electrostatic characteristics stem from the regularly spaced charge pattern of ATDNA major groove and the commensurate helical geometry of homopolymeric ATDNA. As previously shown for the case of cobalt hexamine with RNA,^28^ deeper binding of cations is destructive for mediating inter-helical attraction. Therefore, we reason that the shallow binding of Mg^2+^ plays a critical role in mediating DNA-DNA attraction, as it positions Mg^2+^ ions closer to the opposing helix and facilitates cation-DNA charge correlations.

Our entropy analyses show that DNA condensation incurs significant entropic penalties for both DNA and cations, disfavoring their assembly for both ATDNA and MixDNA sequences. Interestingly, the loss of cation entropy is the largest differentiating factor, creating a strong driving force towards dissociation in the case of MixDNA. ATDNA, however, appears to retain substantial levels of cation entropy, which lowers the barrier to condensation. Nonetheless, cation entropy poses a barrier to assembly for both ATDNA and MixDNA. However solvent entropy overall favors condensation, with ATDNA solvation shell providing a stronger driving force dominated by cation dynamics and electrostatic interactions. Our analysis shows that the contribution of Coulomb energy between DNA pairs provides the driving force. The unique topology of ATDNA with a dynamic two-layer cation shell offers better screening than MixDNA

Specifically, homopolymeric ATDNA line up the partial charges in the major groove in a continuous and periodic fashion, which not only enhances cation binding but confers significant mobility to hydrated Mg^2+^ with shallow binding. This gives rise to a fluidic chain of cations along the DNA major groove rather evenly spaced with the phosphate backbone (which is partially neutralized by electrostatically bound cations). The inter-digitated charge patterns create “zipper-like” charge-charge correlations via orientational coupling. DNA assembly is driven by electrostatic attraction between correlated charges of opposite signs. It is worth noting that, in spite of resembling positional arrangement, our proposed mechanism is physically different from the KL zipper model. First, the KL model starts with cations already bound in the major groove based on empirical observations, while the major-groove cation binding here is substantiated with physical MD simulation. We further show cation binding to be sequence and cation dependent, which is a fundamental difference between ATDNA and MixDNA. Second, the KL model presumes static cation binding in evenly spaced sites that are unphysical, while the cations in the major groove here are observed to retain high levels of mobility, evident from their continuous spatial distributions and entropy calculations. Our studies indicate that the cation dynamics not only make key entropic contributions but also allow exchanges (hops) between adjacent helices. Cation dynamics is thus a novel observation of this study, which requires certain surface features and shallow cation binding.

On the whole, while often deemed to be dominated by the electrostatics of the phosphate backbone, nucleic acids interactions are *de facto* multifaceted and, perhaps more remarkably, nucleic acids prove to be capable of modulating molecular interactions via a variety of surface restructuring strategies. For example, drastic changes can be made by forming multi-stranded structures such as triplex and quadruplex that have been shown to condense in divalent and even mono-valent counterions.^52,53^ Biochemical modifications such as methylation and hydroxylation occur widely in biology and have been shown to promote inter-helical repulsion. Our study here addresses a more subtle perturbation of structure via DNA sequence alone, which nonetheless demonstrates the ability to turn repulsion into an attraction in divalent salts. Such multi-stranded structures, modifications, and repeating sequences are abundant in biology. It is important to recognize the peculiarities of these non-canonical structures to elucidate their functional mechanisms at the molecular level. Furthermore, the versatility of nucleic acids in modulating interactions is expected to be able to assist with the synthesis of functional or therapeutic assemblies such as DNA origami and lipid nanoparticles.

## Acknowledgement

We thank NYUAD for their generous support for the research. This research was carried out on the High-Performance Computing resources at New York University Abu Dhabi. S.K and W. H. would like to thank the REF Grant 317 and AD181 faculty research grant. We thank Richard H. Henchman for his help with the program CodeEntropy.

