## Supporting Information for "Sequence-dependent orientational coupling and electrostatic attraction in cation-mediated DNA-DNA interactions"

### Supporting Information Available

### Molecular modeling of DNA duplexes

The initial structures of both ATDNA and MixDNA constructs were created using Nucleic Acid Builder (NAB),[31] assuming B-form geometry. ATDNA refers to a 20bp duplex consisting of two homopolymeric chains, dA20 and dT20, while MixDNA represents a 20bp duplex with a mixed sequence of GCA TCT GGGC TATA AAA GGG and its complementing sequence. These structures were duplicated to create two parallel arrays, which were then placed in a rectangular cell with the long axis aligned to the z-coordinate. This resulted in an initial simulation box size of $11.8nm \times 11.8nm \times 6.8nm$. Periodic boundary conditions (PBC) were applied to extend the DNA to infinite length. To solvate the DNA, we used explicit water and ions. We studied the ATDNA and MixDNA systems in pure $Mg^{2+}$, as well as ATDNA in two different mixed salt conditions. Table S1) summarizes each simulation setup and salt conditions.

**Table S1**

| Simulation system | ATDNA $Mg^{2+}$ | MixDNA $Mg^{2+}$ | ATDNA $Mg^{2+}/Na^{+}$ | ATDNA $Mg^{2+}/Na^{+}/Spermine^{4+}$ |
|---|---|---|---|---|
| Salt condition | 52.5mM $Mg^{2+}$ | 52.3mM $Mg^{2+}$ | 15.2mM $Mg^{2+}$ 119.8mM $Na^{+}$ | 16.6mM $Mg^{2+}$ 86.5mM $Na^{+}$ 0.21mM $Spermine^{4+}$ |

### General MD simulation set up

All simulations were carried out using GROMACS 2018.5[32] suite of programs. We used amber99sb_parmbsc0[33–36] force field for DNA, Smith and Dang parameters for Na[54], and Cl by Cheatham group[55], and TIP3P[37] for water molecules. For magnesium, we used NBFIX[26] parameters.

Once the simulation system is set up we employed a 5000-step energy minimization to remove any bad contacts resulting from the random placement of water and ions. The minimized simulation set ups later used for MD simulations.

Next, an equilibrium procedure involving volume and solvent was performed on the minimized structures. Specifically, we carried out two nanoseconds of constrained MD at NPT to adjust the simulation volume, followed by 200-ns of constrained MD at canonical ensemble to equilibrate the water and ions around the restrained DNA molecules.

The equations of motion were integrated using the Leap-Frog scheme with a 2-fs time step. Particle mesh Ewald (PME) summation method was used to treat electrostatics, with a grid spacing of 0.12 nm and an interpolation of order 4. Non-bonded interactions and neighbor searches were cutoff at 11 Å, and the list was updated every 40 steps. The covalent bonds of water and nucleic acid were constrained using SETTLE[56] and LINCS[57] algorithms, respectively. We used Berendsen thermostat[58] and Parrinello-Rahman barostat[59] for NPT simulations and velocity scaling[60] for NVT simulations. During the equilibration, all heavy atoms of DNA were restrained with a stiffness constant of 1000 kJ $mol^{-1}$ $nm^{-2}$, while water and ions were allowed to move freely. The output coordinate of the NVT simulation was used as the starting point for the following Well-tempered metadynamics simulations.

### Well-tempered metadynamics simulations

Well-tempered Metadynamics (WTMD)[30] was employed to extensively sample the conformational space of two parallel DNA pairs. To describe the dynamics, we employed two collective variables (CVs): inter-helical distance ($d$) and inter-helical rotation ($\theta$) (Fig.1a-b). During the WTMD runs, we constrained the orientation of the center helix (H1) by applying a harmonic restraint with a stiffness constant of 1000 $kJ \cdot mol^{-1} \cdot nm^{-2}$. Gaussians were initialized with a magnitude of W=0.6 kJ/mol and deposited every 1 ps with a gradually decreasing bias factor of 6.0. The widths of the Gaussians were set to $\sigma_d = 0.1$ nm for $d$ and $\sigma_\theta = 0.2$ rad for $\theta$, respectively. The heights of the deposited Gaussians and the collective variables were monitored during the simulations (Fig.1c). Following,[41,61] the decay of the heights of the deposited Gaussians to $< 0.005$ kJ/mol was used as a threshold to assess the convergence. In addition, the convergence of each simulation was assessed by performing block analysis (Fig. S3).

### Data Analysis

We monitored the position of each cation in time. This data is then used to compute the average density and charge profiles, and entropy.

#### Computing the entropy of $Mg^{2+}$ and water using 2PT method

We used the DoSPT program[44] to apply the 2PT method and estimate the entropic contribution of $Mg^{2+}$ with its hexa-hydrated shell (i.e., $Mg(H_2O)_6{}^{2+}$). We studied cation entropy in the condensed state (with inter-helical distances of ~2.8 nm) and in the free state (with inter-helical distances of ~4.0 nm). To do this we first generate a pool of structures sampling each state. For that we performed 150-ns-long MD simulations at 300 K, with the inter-helical distances restrained to the states defined. We selected a conformation every 5 ns (resulting in 30 starting points). From each conformation selected we conducted a 120-ps-long NVT simulation with $\delta t = 1$ fs resolution and we recorded the forces, velocities, and coordinates in 4 fs time intervals. The saved trajectories were then used in DoSPT program to estimate the entropy. The convergence of the entropy calculation was assessed using trajectories of varying lengths (80, 100, 120, and 150 ps), as shown in Fig. S4. The error bar was computed as the standard deviation by taking the average of these 30 independent DoSPT runs, it accounts for any variability in the results that could arise from small differences in the initial conditions or other factors.

Similarly, we estimate the entropy of solvent water using the same approach. The 2PT calculation of water entropy is known to converge quickly, short (~10 ps) MD simulations provide very accurate values of energy and entropy,[46] we run 20ps in this study. We evaluate the bound water defined by the R1 hydration shell (Fig. 3c) and bulk water, respectively. A

~50 mM $MgCl_2$ solution was simulated for 50ns as pure-solvent system in order to compute the entropy of water in bulk ($S_{bulk}$). As for the entropy of bound water, we pick the same starting points applied in entropy calculation of $Mg^{2+}$. This time only the free state is considered, the water molecules within 3.5Å (R1 shell) of DNAs' surface were used to perform DoSPT calculations ($S_{bound}$). The entropic contribution of bound water released to the bulk ($\Delta n$) occurred during free-state-to-condensed-state transition is was then expressed as $\Delta S_{water} = \Delta n(S_{bulk} - S_{bound})$ (Table 1).

#### Computing the entropy of DNA duplexes using multiscale cell correlation method

Using the CodeEntropy program, which implements MCC theory,[42,43] we calculated the macromolecular entropy of DNAs. Similar to the cation entropy analysis, we utilized the two end-state trajectories. We performed 100-ns-long MD simulations at canonical ensemble with a temperature of 300K and restrained the inter-helical distance at the desired values to record the forces every 10 ps. These forces were loaded into the MCC program to estimate the macromolecular entropy. To estimate the error, we divided the trajectory into two equal parts. In our study, we found that the topographical entropy $S_M^{topo}$ was small due to the relatively stable backbone structure and was hence neglected.

#### Cation density profiles and water structuring

We compute the average cation density profiles and tetrahedral order parameters from the 150 ns long MD trajectories described above. For density analysis, we investigate (e.g. pure $Mg^{2+}$, $Mg^{2+}/Na^{+}$ and $Mg^{2+}/Na^{+}/Spermine^{4+}$) at the condensed and free states. Due to linear charge distribution of $Spermine^{4+}$, we divided the molecule into 4 parts according to the position of its four nitrogen atoms during the calculation (Fig. 7c, bottom).

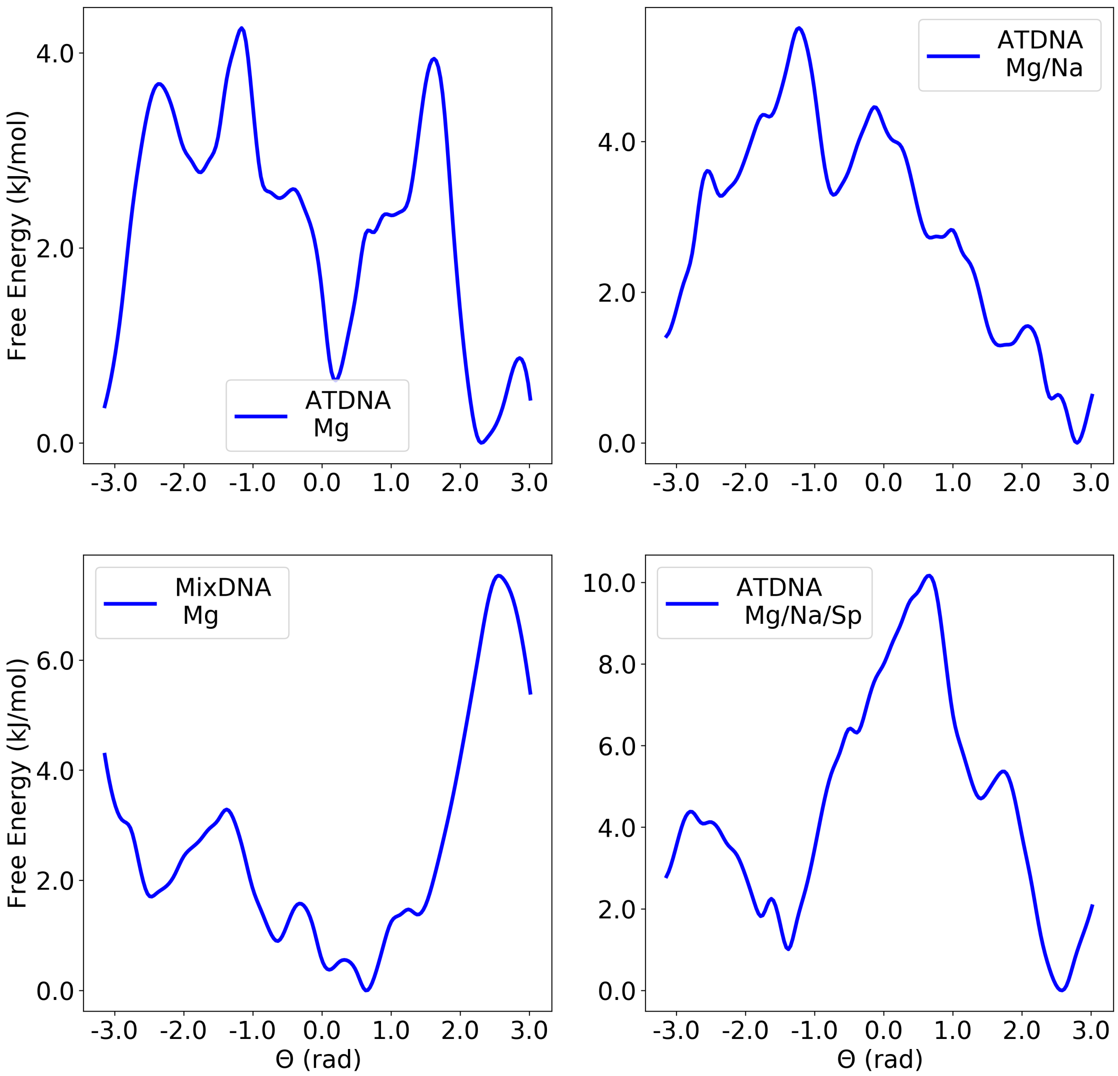


**Figure S1: Free energy profiles projected on the collective variable $\theta$.**

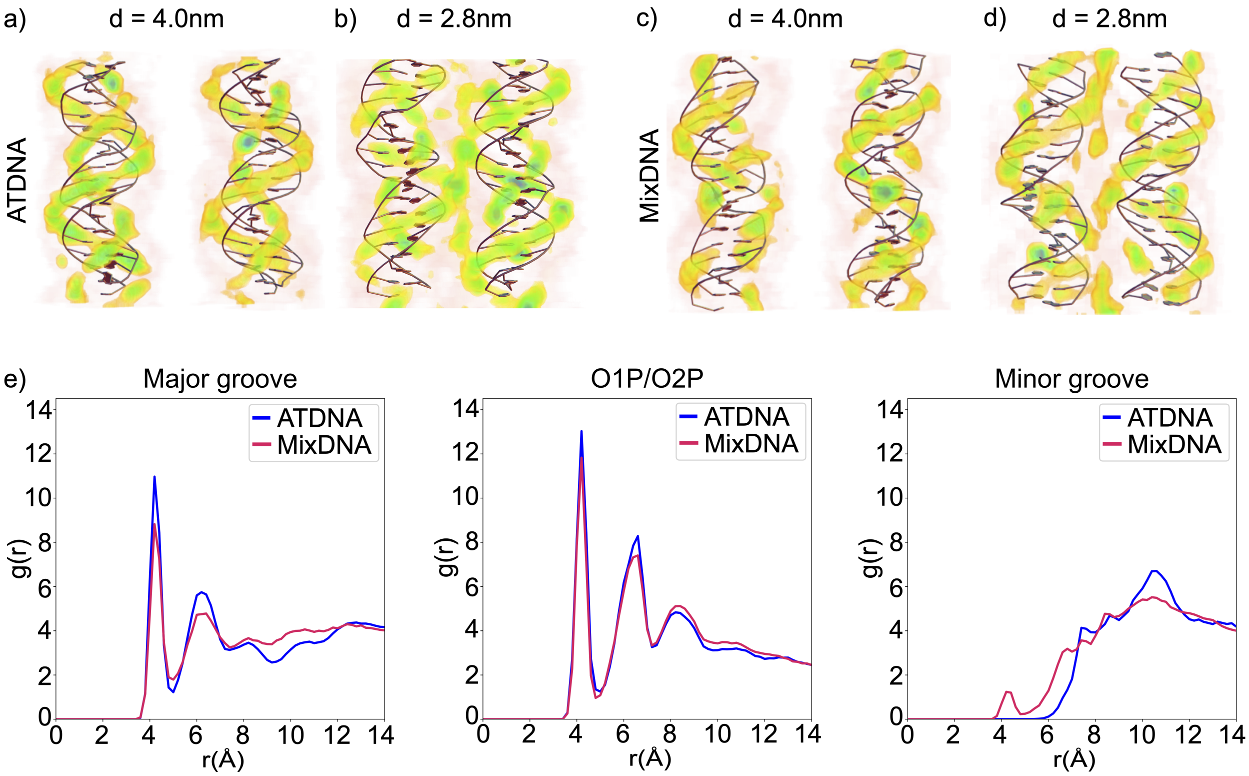


**Figure S2: The change of average $Mg^{2+}$ ion distribution around the duplexes during the dissociation-to-association transition.** We plot the 3D ion density map for a) free state and b) condensed state of ATDNA and the c-d) two sates for MixDNA. e) Radial distribution function of $Mg^{2+}$ ions from the major group atoms (left), phosphate backbone (central) and minor groove atoms (right) for ATDNA (blue solid) and MixDNA (green solid) in pure $Mg^{2+}$ solution, respectively.

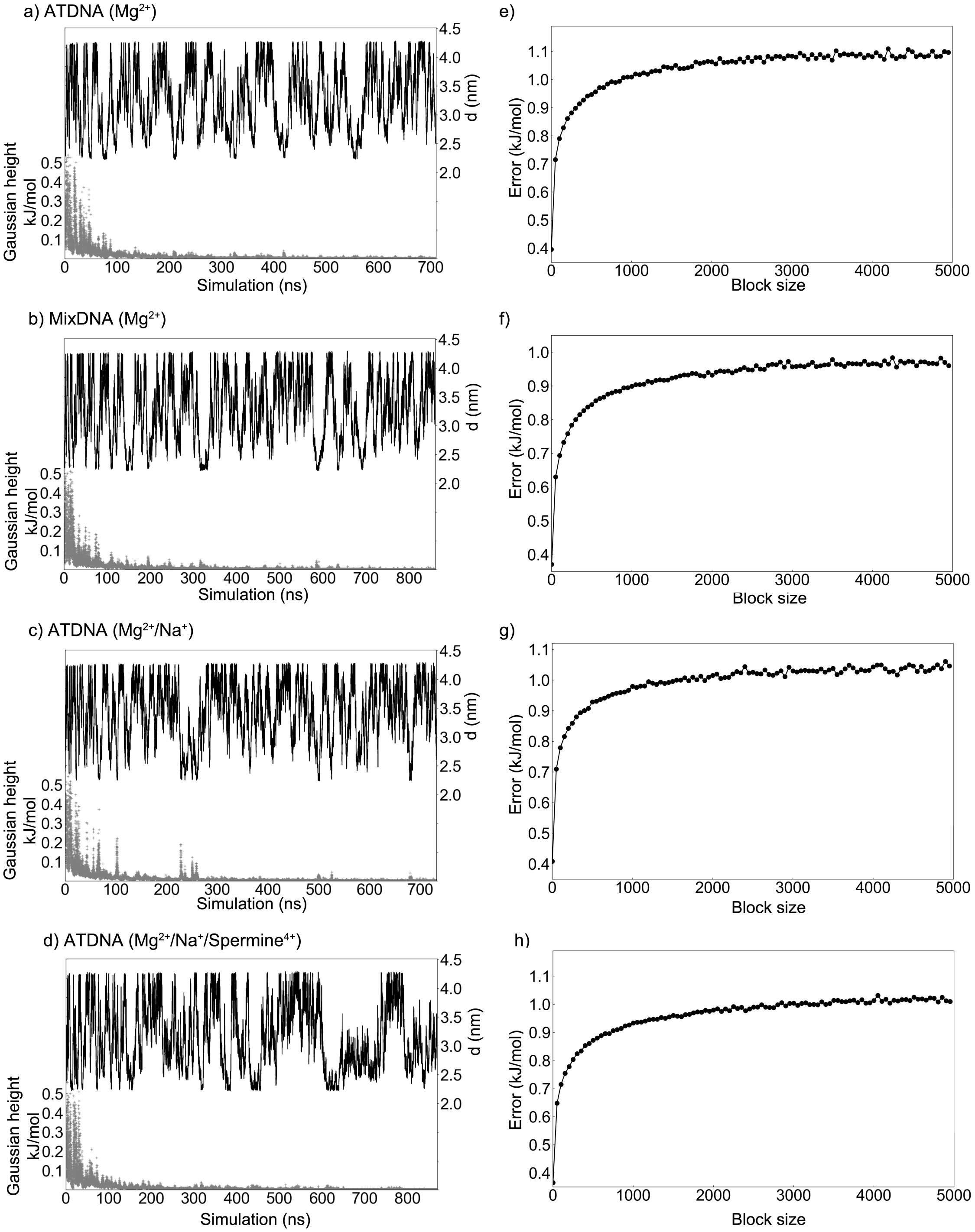


**Figure S3: Time evolution of the collective variable d (black) and Gaussian hills (gray) during metadynamics simulations and block analysis.** a)/e) ATDNA in $Mg^{2+}$, b)/f) MixDNA in $Mg^{2+}$, c)/g) ATDNA in $Mg^{2+}/Na^{+}$ and d)/h) ATDNA in $Mg^{2+}/Na^{+}$/Spermine$^{4+}$.

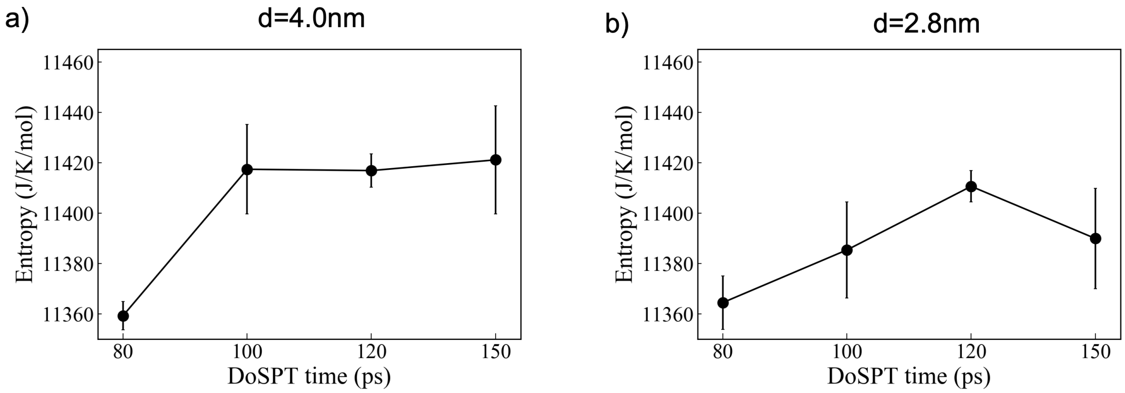


**Figure S4: Convergence of the DoSPT entropy calculations for the ATDNA/$Mg^{2+}$ system. The graph illustrates the convergence behavior of the DoSPT calculations as a function of simulation time.**

**Movie 1. Molecular simulation of ATDNA in $Mg^{+2}$**

**Movie 2. Molecular simulation of ATDNA in a mixture of $Mg^{+2}$, $Na^{+}$ and $Sp^{+4}$**

### Graphical TOC Entry

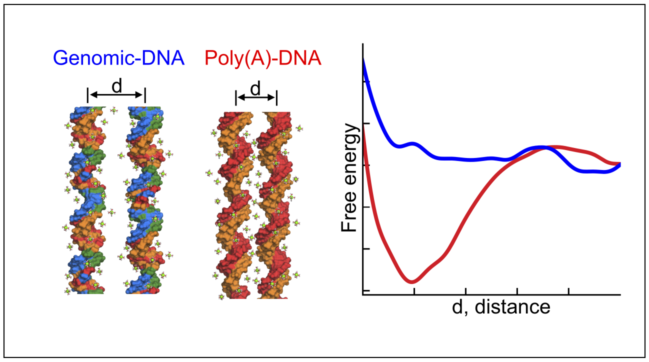
